# Independent evolution towards large body size in the distinctive Faroe Island mice

**DOI:** 10.1101/747097

**Authors:** Ricardo Wilches, William H. Beluch, Ellen McConnell, Diethard Tautz, Yingguang Frank Chan

## Abstract

Most traits in nature involve the collective action of many genes. Traits that evolve repeatedly are particularly revealing about how selection may act on traits. In mice, large body size has evolved repeatedly on islands and under artificial selection in the laboratory. Identifying the loci and genes involved in this process may shed light on the evolution of complex, polygenic traits. Here, we have mapped the genetic basis of body size variation by making a genetic cross between mice from the Faroe Islands, which are among the largest and most distinctive populations of mice in the world, and a laboratory mouse strain selected for small body size, SM/J. Using this F2 intercross of 841 animals, we have identified 102 loci controlling various aspects of body size, weight and growth hormone levels. By comparing against other studies, including the use of a joint meta-analysis, we found that the loci involved in the evolution of large size in the Faroese mice were largely independent from those of a different island population or other laboratory strains. We conclude that colonization bottleneck, historical hybridization, or the redundancy between multiple loci have resulted in the Faroese mice achieving an outwardly similar phenotype through a distinct evolutionary path.

## Introduction

Discovering the genetic basis of naturally occurring variation is an important step towards understanding how traits change through time. Knowing which genes are involved in controlling traits may allow, among other things, inferring how selection operates in shaping organisms and explaining natural phenomena like rapid evolution (Barton and Keightley, 2002). Selection is a deterministic process, which means that unlike neutral processes like drift, a hallmark of selection is that similar phenotypic outcomes can and do evolve independently. This phenomenon, also known as parallel evolution, is often seen as indictive of selection (Schluter *et al*., 2004). By studying parallel evolving systems, we may gain insight into the relative roles of selection and chance (Conte *et al*., 2015). Despite many remarkable studies in identifying the genes associated with parallel changes (Chan *et al*., 2010; Conte *et al*., 2015; Zhen *et al*., 2012; Meyer *et al*., 2018), most tend to focus on traits controlled by only a handful of genes. Fewer still connect what occurred in nature to results obtained in the laboratory. We therefore do not know if the same rules of parallelism or convergence hold for more complex traits like body size and weight (but see Chan *et al*., 2012).

Island populations of mice, in this sense, represent an outstanding opportunity to study parallel evolution: the house mouse is exceedingly successful in colonizing diverse habitats, including numerous remote islands, many at high latitudes. Following colonization, they have often evolved to large body size (Foster, 1964; Lomolino, 1985). Such events, repeated over and over again, represent replicated natural experiments. There are many parallels to changes in body size and weight in laboratory mice, which have been the subject of many classical studies in quantitative genetics, developmental genetics and physiology (MacArthur, 1944; Bünger *et al*., 2001; Renne *et al*., 2006; Cheverud, 1996; Chan *et al*., 2012). Together, these resources present a unique opportunity to study the genetic basis of evolutionary change in nature, and to make connections to results obtained among laboratory strains.

We chose to study the mouse population on the Faroe Islands, because of their body size and as a (extreme) representative of the mice in the North Atlantic. The Faroe Islands are an island group of 18 major islands in the North Atlantic, six of which are inhabited by house mice. Since their discovery more than a century ago, the Faroese house mice have attracted considerable interest, on account of its large body size and distinctive morphology (Eagle Clarke, 1904). So remarkable were the Faroese mice that they have variously been classified as the new subspecies *faeroensis*, or even *mykinessiensis*, from a single island Mykines of only 10 km^2^ in area (Eagle Clarke, 1904; Degerbøl, 1942; Berry *et al*., 1978). To date, the population on Gough Island represents the most extensive studied thus far in the context of island gigantism in house mouse (Gray *et al*., 2015; Parmenter *et al*., 2016). Unlike the mid-Atlantic Gough Island population, whose continental source population remains unclear (Gray *et al*., 2014), the Faroese mice show close links to other Northern Atlantic populations in morphology and genetics (Berry *et al*., 1978; Jones *et al*., 2011; Jones *et al*., 2012). In addition, our genomic survey has revealed a historical hybridization event that temporally coincided with Viking movements and the Danish takeover of the Faroe Islands (Jones *et al*. in prep.). Specifically for genes controlling body-weight, we have previously detected evidence of selective sweeps in the Faroese mice at two loci found to be repeated involved among laboratory mice selected for body weight changes (Chan *et al*., 2012). This makes the Faroese mice a compelling example of island gigantism for a large-scale genetic mapping study.

## Results

To enable direct comparisons between laboratory and wild mice in their genetic basis of body size and weight, we organized a field expedition to collect live mice from the Faroe Islands. We focused on Mykines (“MYK”), the westernmost major Faroese island, because its mouse population were consistently the largest and most distinctive in the Faroes (Eagle Clarke, 1904; Degerbøl, 1942; Berry *et al*., 1978). For the purposes of line crosses, the MYK population also were most inbred (Jones *et al*., 2011; Jones *et al*. in prep.), making it likely that sampled individuals would be representative of the population at large. Following a generation of breeding under common conditions, we set up crosses with the small SM/J strain to determine firstly the genetic basis of body weight and length variation using a quantitative trait locus (QTL) mapping approach, and additionally to assess the extent of QTL sharing with other examples of body weight evolution, principally by a combined QTL meta-analysis using previously published QTL studies involving the SM/J strain, mostly against the large LG/J line (Cheverud, 1996; Stylianou *et al*., 2006; Parker *et al*., 2011).

Overall, the MYK mice were significantly larger and heavier than SM/J mice between weeks 8 to 13 of age (Fig. 1B; MYK vs. SM/J, length at 13 weeks: 9.2 vs. 8.2 cm; Student’s *t*-test, *h_1_* > *h_0_*; males, 13 weeks: *t* = 6.7, df = 13.6, *P* < 5.8 × 10^-6^; weight at 8 weeks, males: 19.5 vs. 16.0 g: *t* = 3.18, df = 12, *P* < 0.004; females: 17.8 vs. 16, *t* = 1.6, df = 6, *P* = 0.08, non-significant; 13 weeks, males: 22.5 vs. 18.9 g, *t =* 3.1, df = 6, *P* < 0.004; females: 20.5 vs. 18.1 g, *t* = 1.5, df = 11, *P* = 0.08, non-significant). From 12 F1 full-sib families (MYK x SM/J in both directions), a total of 841 F2 mice were born (Fig. S1). For each F2 mouse, we collected a battery of measurements over 16 weeks of age (Table S2). The set of measured traits included 17 length, weight and growth hormone measurements. Males were on average heavier than females, weighing 15.9 – 22.4 g from week 4 to 16 vs. 14.7 – 19 g in females. We observed a slight, but significant parent-of-origin effect, but only in females: female mice having a MYK paternal grandmother tended to be born lighter, but gained more weight through to adulthood (Fig. S2; N = 4242 observations in 606 individuals, repeated measures ANOVA with time and cross direction; effect for cross direction over time: *P* < 0.89 in males and 0.0003 in females). In an F2 intercross design, all F2 females but not males consistently carry the paternal grandmother X chromosome, this effect was consistent with the MYK X chromosome being associated with lighter birth weights but greater growth rate over 16 weeks.

**Figure 1.**
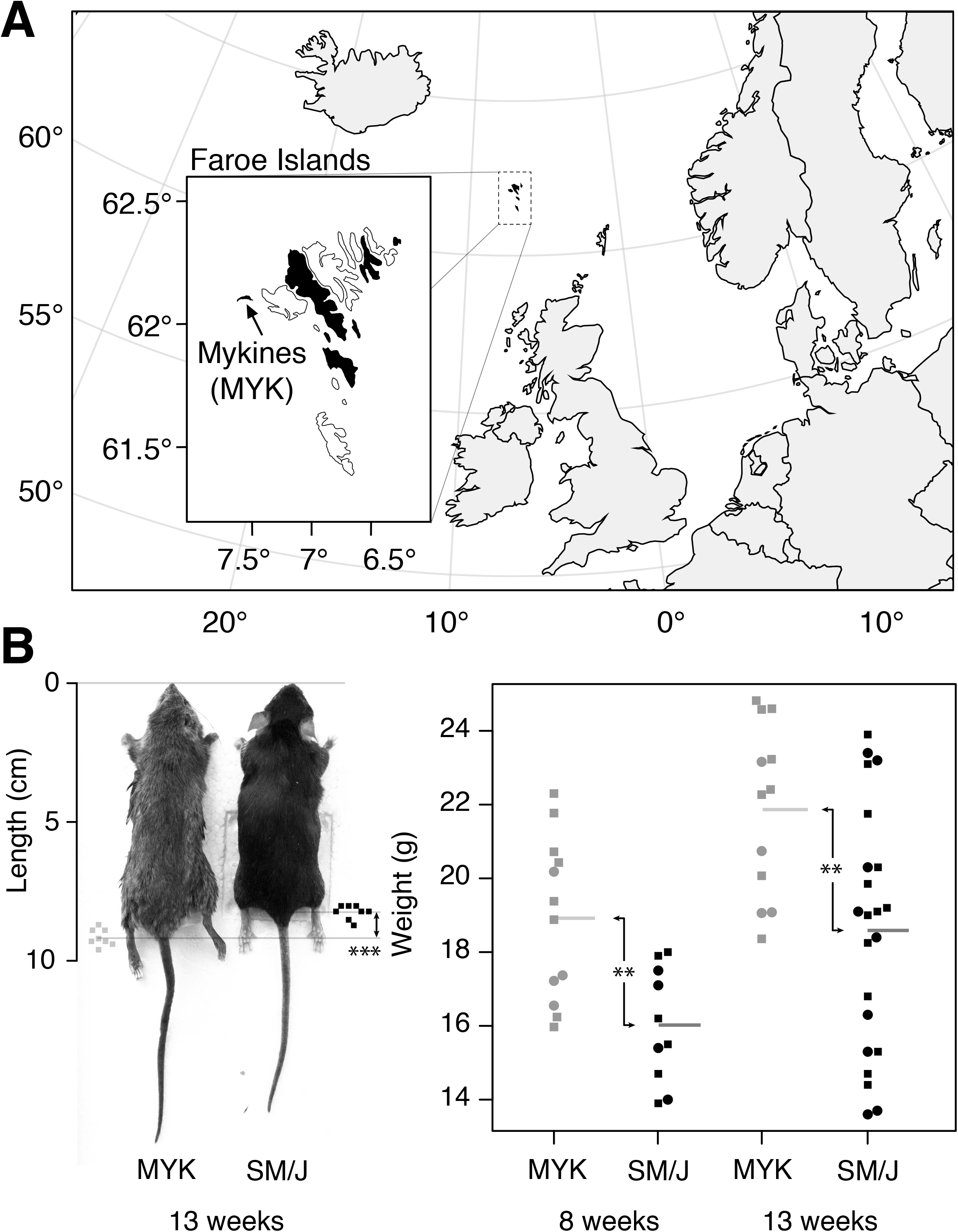
The large-bodied Faroese house mouse. The Faroe Islands are an island group of 18 major islands in the North Atlantic (inset), six of which are home to wild house mice (*Mus musculus*, mainly of *domesticus* subspecies, historically *faeroensis*; black) (Berry *et al*., 1978). Mykines (MYK), the westernmost island measuring 10 km^2^, is home to a distinctive mouse population (“*mykinessiensis*”). These mice are among the largest wild mice in the Faroe Islands and the world. A genetic cross was set up between the MYK mice (grey) and the small laboratory strain SM/J (black) to investigate firstly the genetic basis of weight and length variation in the Faroese mice, and secondly the extent of sharing in the genetic loci underlying body weight changes between laboratory and wild mice. Squares: males; circles: females; **: *P <* 0.004; ***: *P <* 1×10^-5^.

In order to perform quantitative trait locus (QTL) mapping to find associations between the genotype and phenotype data, we sequenced the parental lines and constructed a genetic map using genotypes from a two-enzyme version of restriction-site associated DNA sequencing (RAD-seq; Poland *et al*., 2012; Witte *et al*., 2015).

We approached the genetic mapping in three steps. First, we mapped each trait singly. Then we focused on the underlying structure of the data by either fitting growth curves over the entire growth series in each individual, or by extracting the major axes of variation in the dataset via principal components analysis. Finally, we performed a composite joint mapping by integrating data from additional QTL datasets involving SM/J, in order to examine the extent of QTL sharing between laboratory mice and MYK mice, as an example of island gigantism in the wild.

Overall, our mapping revealed a strong genetic basis for trait variation in this cross. For all but one of the 17 measured traits, we found 2 to 8 QTLs that together explained on average 24% of the variance in a trait (Table 1; median: 5 QTLs per trait, explaining on average 4.9%; combined, the QTLs explain 8 – 45.9% of variance in a given trait).

**Table 1.**
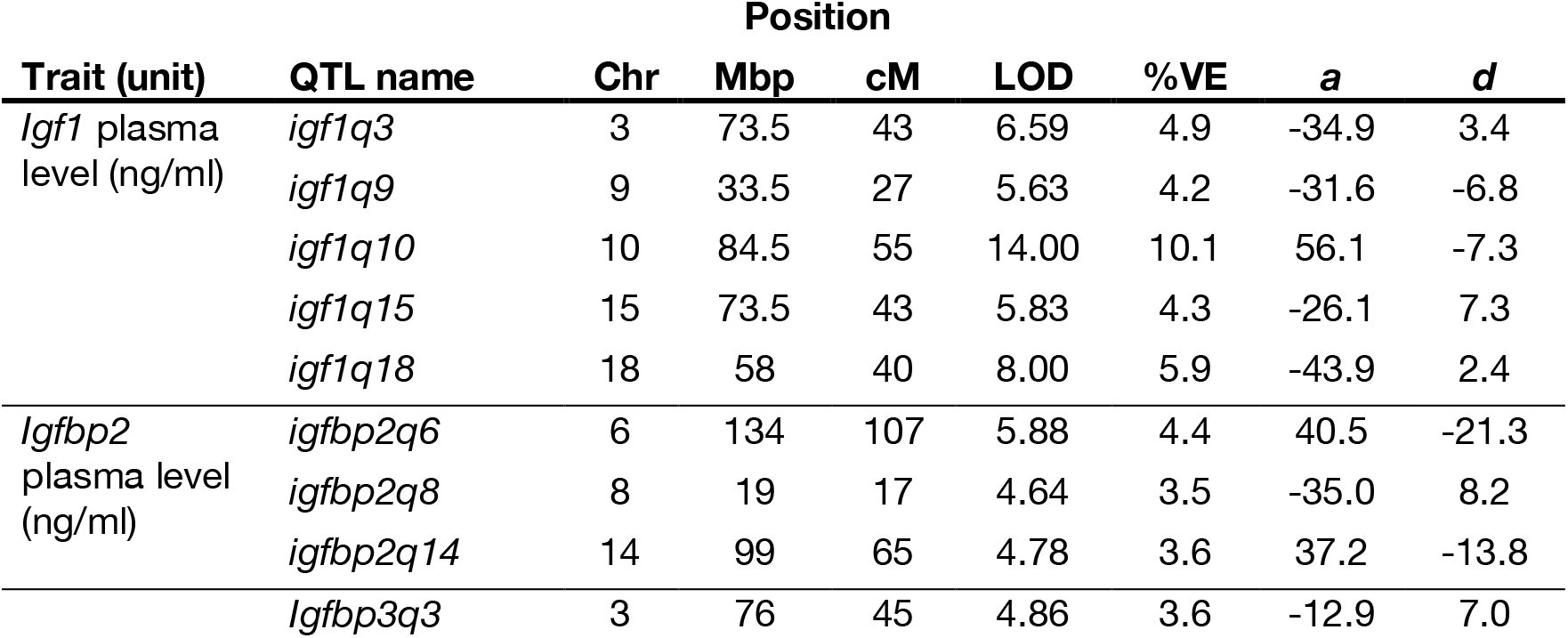
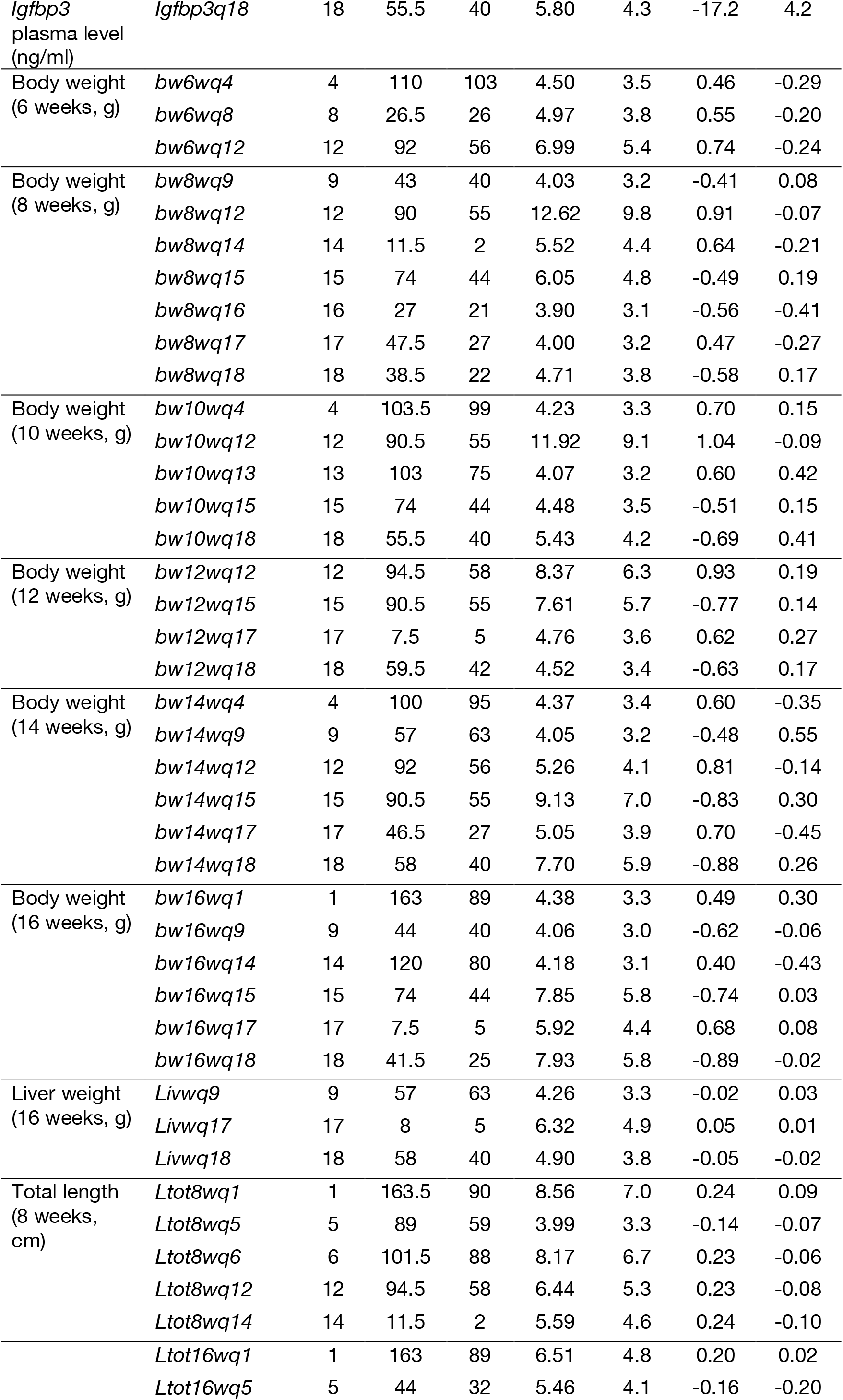
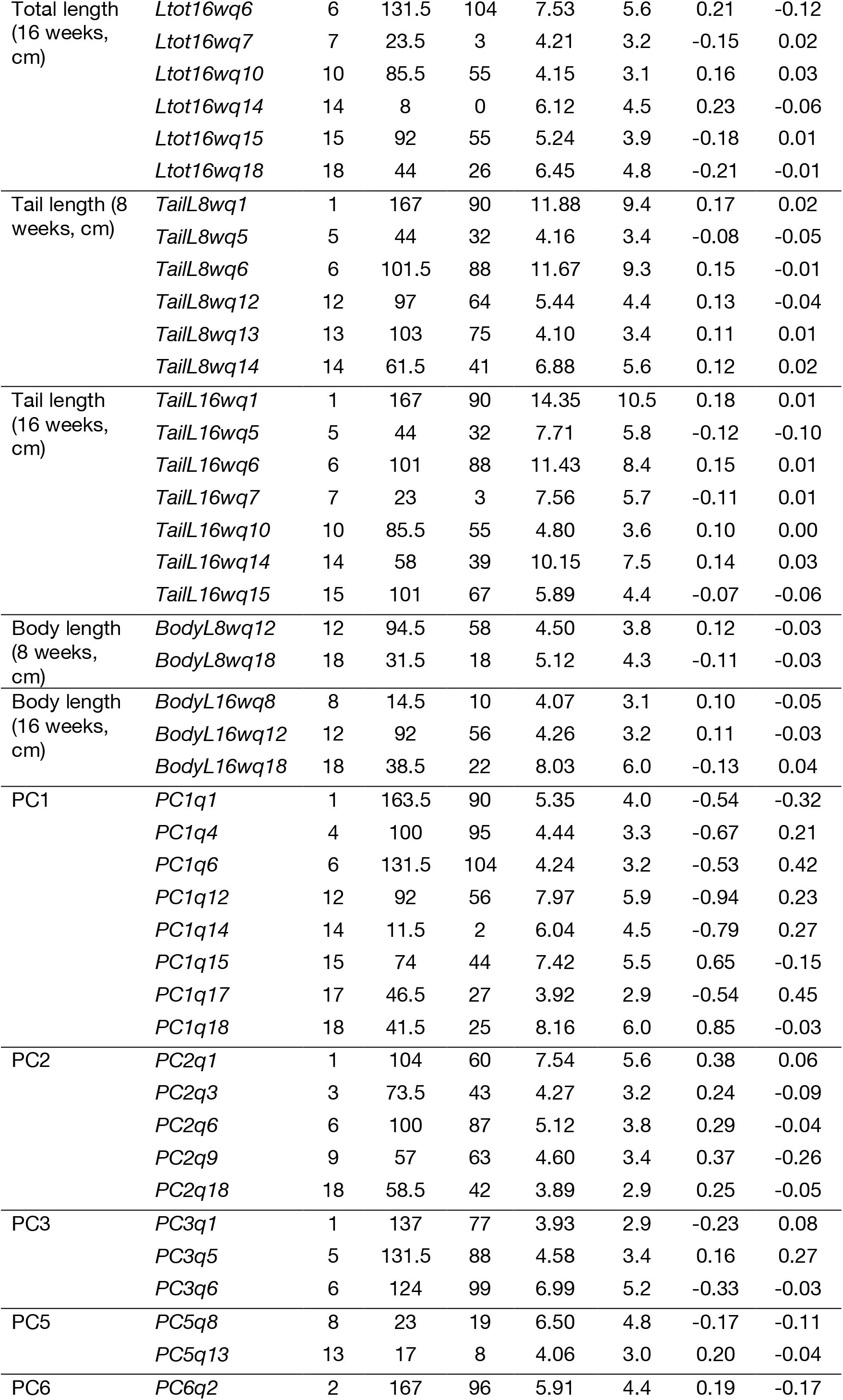
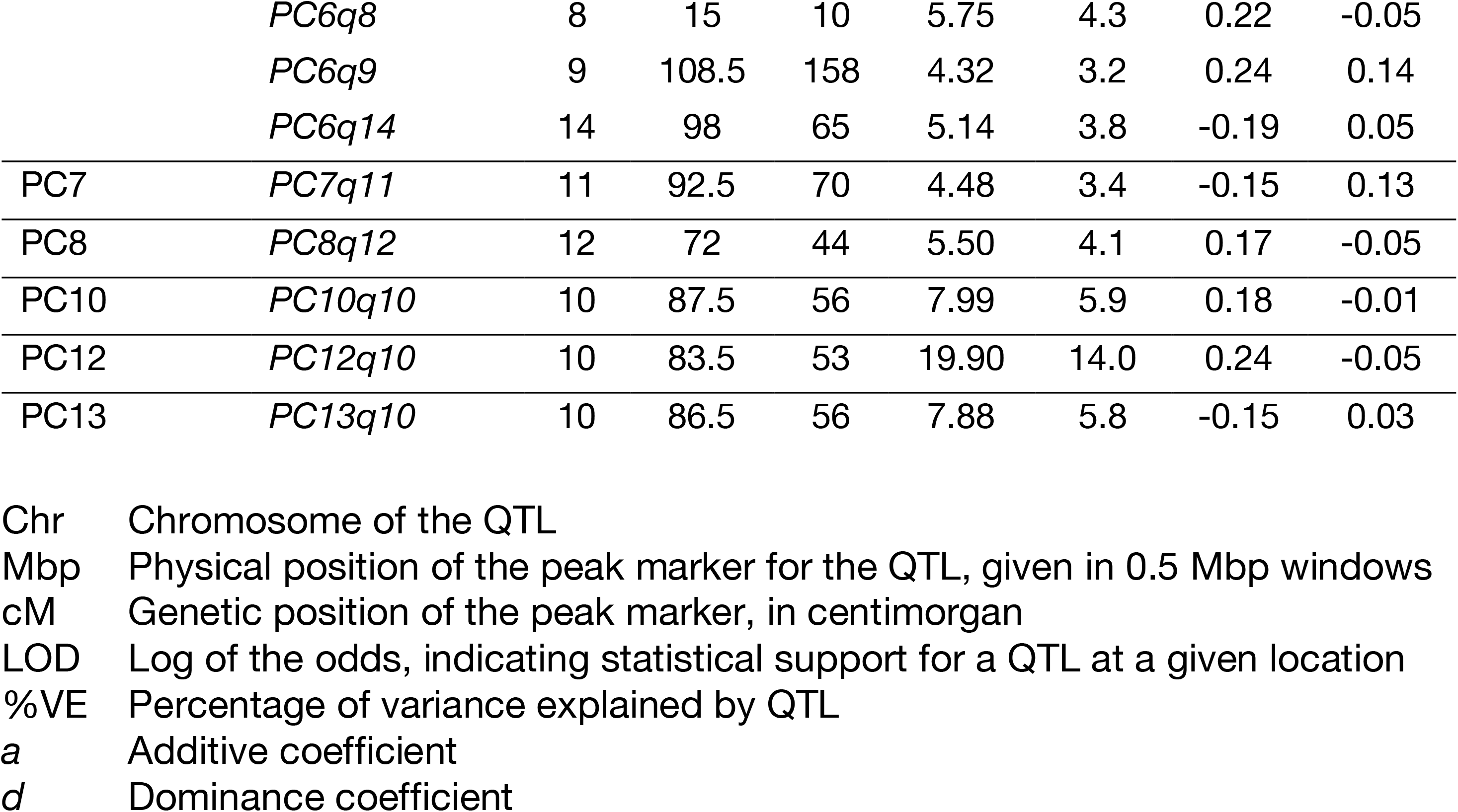
QTL locations and effect sizes.

Among the strongest QTL is a locus on Chr10 (84.5 Mbp), *igf1q10*, that controls the blood plasma level of the growth hormone *Insulin-like growth factor 1* (*Igf1*, LOD: 14.0, 10% variance explained). This QTL overlaps with the *Igf1* gene itself (Chr10: 87.9 Mbp) and a previously reported QTL *Igf1q4* involving SM/J and another laboratory strain MRL/MpJ (Leduc *et al*., 2010). Here, we can recover the same 5’UTR C/A variant rs29342496, with the MYK allele also carrying the A variant, each of which we estimated to increase *igf1* level by 56 ng/ml (cf. an average of 391 ng/ml among F2 mice). This is largely consistent with the contrast seen in crosses between laboratory strains (Rosen *et al*., 2000; Leduc *et al*., 2010). Like those previous studies, we observe no protein coding changes at *Igf1* that might affect protein degradation. Taken together, our data suggests that SM/J may carry a regulatory variant affecting the circulating *igf1* protein level.

The strongest QTL in this dataset, *TailL16wq1*, is located on Chr1 (167 Mbp) and controls variation in tail length (16 weeks, LOD: 14.4, 10.5% variance explained). This locus—or other tightly linked ones—also controls tail length at 8 weeks, and body weight at 16 weeks. Unlike *igf1q10* on Chr10 for plasma levels of *igf1*, this is a morphological trait, not a gene-specific one. Close examinations of the confidence interval (Chr1:161.5 – 169.5 Mbp) revealed 63 protein-coding genes, including the genes *Suco* and *Lmx1a*, whose knockout phenotypes show bone ossification and tail phenotypes, as well as multiple classical spontaneous short tail mutants for *Lmx1a* (Sohaskey *et al*., 2010; Wahlsten *et al*., 1983). Overall, our data suggests that the tail of a mouse may be governed by a smaller set of loci, because for this trait we could detect three major QTLs with >10 LOD support, with 9 total QTLs (6 shared) between the 8- and the 16-week measurements. In fact, at week 16 of age, the seven QTLs could together explain nearly half of the variance.

**Fig. 2.**
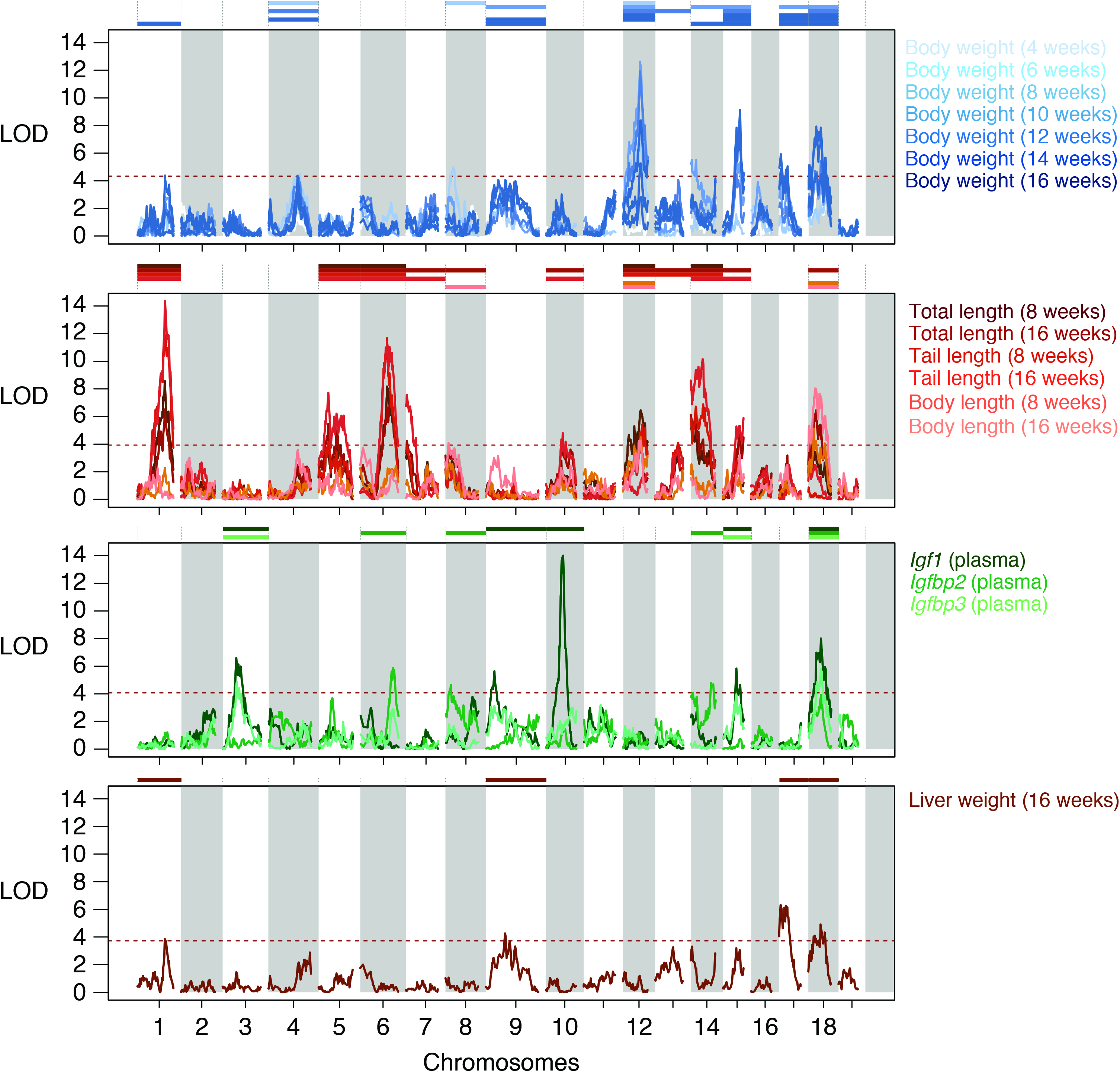
A polygenic architecture in body length and weight variation. Linkage mapping was performed to detect loci affecting body weight (blue); lengths (red); growth hormone levels (green) and organ weight (brown; x-axis: chromosomes; y-axis: log of the odds statistical support for a quantitative trait locus or QTL affecting a trait). In total, we detected X QTLs at the genome-wide significant threshold of ∼3.9 (red dotted line) on 17 out of 19 autosomes (QTL: coloured bars over the chromosome).

We next turn to body weight measurements. As expected, biweekly weight measurements are highly correlated (R = 0.32, 6 time points, N = 606; Fig. 3A; Table S3). We thus observe that many single trait QTLs overlap, with genome-wide significant QTLs on Chr12, 15, 17 and 18 at multiple time-points. Except for the lack of genome-wide significant QTLs at week 4, there is not a clear trend between the number of QTLs (range: 3 – 7) and the timing of the measurement. Across all growth QTLs, the *bw8wq12* at Chr12: 90 Mbp (significant for 6 – 14 weeks, 2-LOD interval: 89.5 – 94.5 Mbp) has the highest LOD, peaking at 12.6 in week 8. This QTL covers only 7 genes, including *Thyroid stimulating hormone receptor* (*Tshr*), a compelling candidate that plays a central role in metabolism and growth regulation principally though the pituitary–hypothalamus axis. At *Tshr* we also did not find any non-synonymous mutations. The report of maternal effect in a spontaneous mutant may also be consistent with the observed decrease in the effect of this QTL as the mouse ages (Dionne *et al*. 2014). This same gene also showed a strong selective sweep signature in domesticated chicken breeds (Rubin *et al*., 2010). In that study, the authors hypothesized that it was presumably associated with altered photoperiods. Assuming conserved gene functions, that interpretation would suggest a connection to the northerly latitudes of the Faroe Islands.

**Fig. 3.**
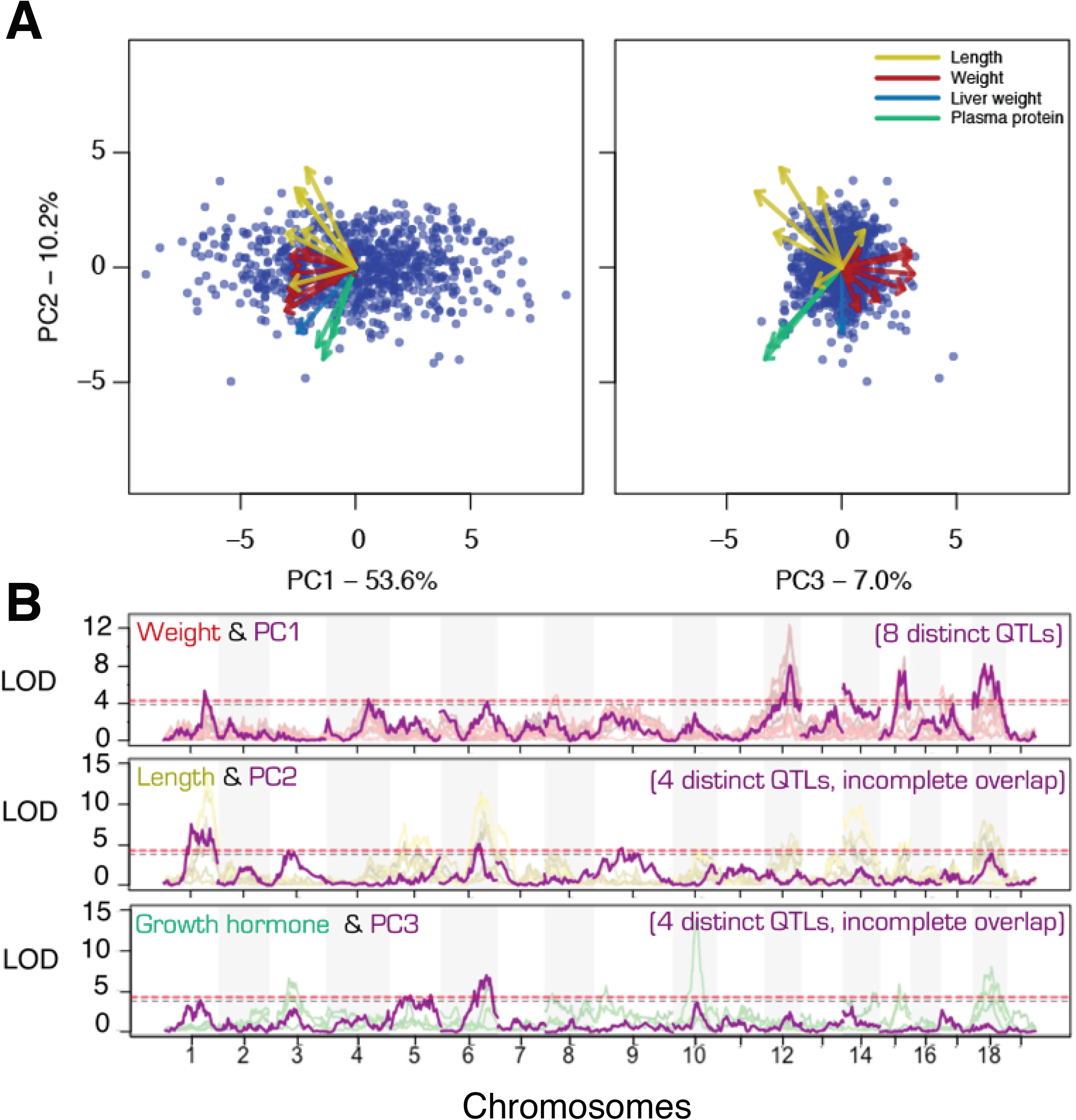
Major modes of variation in body weight and length summarize and recapitulate trait types. A) Individual measurements were resolved into independent modes of variation using a principal components analysis (PCA). There is a strong correlation across all traits, as indicated by the shared direction of each of the trait vectors in both PC1 – 2 and PC 2 – 3 projections (each dot is an F2 individual. Trait vectors are coloured according to trait types; see also Table S3). B) Mapping of the PCs show LOD profiles that largely track with the individual trait types: PC1 with weight, PC2 with length and PC3 with growth hormones.

Due to the broader confidence intervals at the other major QTLs covering a far greater number of genes and the difficulty in assigning gene candidates specifically related to body weight and/or metabolism changes, as opposed to a generic by-product of other functions, e.g., reported weight loss due to mobility issues resulting from the loss of legs, we will turn to mapping and interpreting broader growth curves and major variation modes.

In addition to individual measurements, we also summarized the growth series by fitting Gompertz growth functions (Gompertz, 1825), namely into two parameters: the maximal growth rate *µ* and the final weight or asymptote *A*. The obvious advantage of fitting growth curve is to capture the growth dynamics in an individual animal. However, the assumptions and errors associated with growth curves fitting may also obscure potentially important observations, such as large variations in weaning weight at 4 weeks or drops in body weight over time. Here, we were able to estimate growth parameters for *µ* in 229 and for *A* in 216 individuals, and we detected no QTL at the genome-wide significant level, and a single suggestive QTL for weight *A* at Chr18: 58 Mbp that overlaps with that of the body weight QTLs from weeks 10 – 16. Notably, this QTL also overlaps *igf1q18* and *igfbp3q18*, underscoring the functional involvement of the IGF1 pathway in later—in mouse, post-pubertal—growth (Styne, 2003; Stratikopoulos *et al*., 2008; Courtland *et al*., 2011).

Another way to summarize QTLs would be to extract the major axes of variation within the trait data using principal component analysis (PCA), and map each as a composite trait. In effect, we use PC traits as a way to isolate and map mutually independent growth modes. Visualization of the top two principal components (PC1 and 2, accounting for 54% and 10% respectively) shows that all the measured traits are strongly correlated with each other, with their eigenvectors all loading negatively on PC1 (Fig. 3A; Table S3). Beyond the first level, the different types of traits quickly split into independent directions, such that the PCs 2 and 3 quite effectively summarizes the contrast between length, and liver weight and plasma protein levels; and weight vs. others.

Mapping of these PCs both recapitulates the major QTLs, but also revealed additional loci not discovered in single trait analysis, e.g., *PC2q1* at Chr1: 104 Mbp and *PC3q5* at Chr5: 132 Mbp. More broadly, PC1 nicely summarizes the body weight variation (Fig. 3 and Fig. 4). In the case of the 8 genome-wide significant PC1 QTLs (and two additional suggestive QTLs), they show a classical polygenic architecture associated with body weight and growth.

**Fig. 4.**
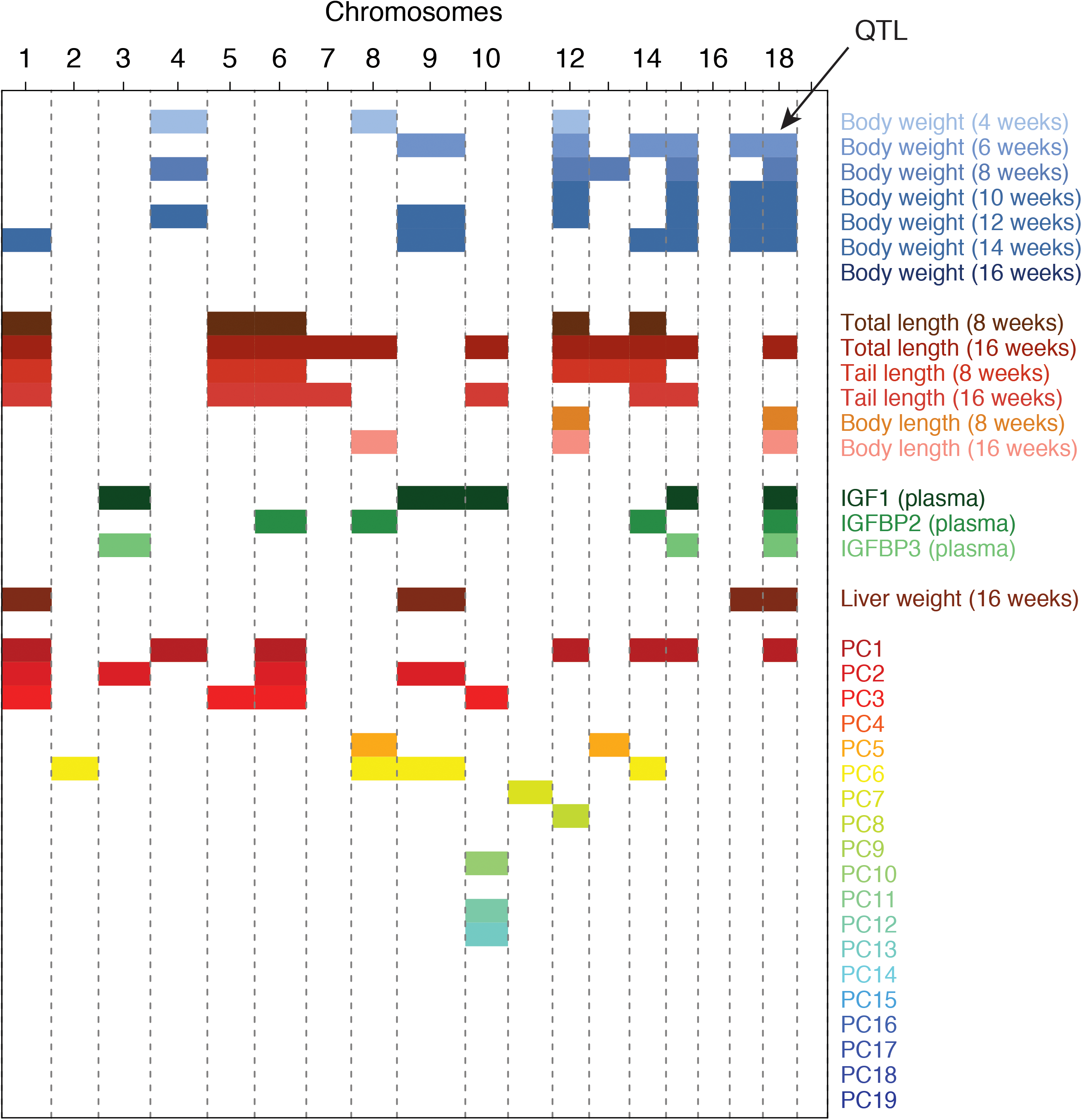
The genetic architecture of body weight and size is polygenic and shows little clustering. A summary representation of all of the significant QTLs recovered in the current study shows that while there is a strong auto-correlation and sharing within a given trait type (weight, size, etc.), the set of chromosomes carrying genome-wide significant QTLs appear to be largely distinct from one type of traits to another. This also result in the absence of a single or few “super-gene” clusters who account for much of the observed variation in the trait.

A global comparison of the chromosomes carrying QTLs across the different trait types also recapitulates the PCA shown in Fig. 3. With the possible exception of *igf1* level, most types of traits are controlled by many loci scattered across the genome and tend not to cluster into single major-effect chromosomes.

We next asked if there was evidence of QTL sharing in examples of parallel body weight changes in laboratory and wild mice. To do so we collected data from other laboratory QTL crosses (“panels”) involving the SM/J strain, namely the LG/JxSM/J crosses by the Cheverud and later the Palmer laboratories and the NZB/BINJxSM/J cross by Stylianou et al. 2006, for a total of 4552 mice (Table S4) (Cheverud, 1996; Stylianou *et al*., 2006; Norgard *et al*., 2008; Parker *et al*., 2011). An inherent limitation in many meta-analysis is the variation in protocols and the underlying datasets. We found that among our measurements, 16-week weight was the only trait that was similar enough across the datasets to allow a meta-analysis. We merged the SM/J-polarized genotypes from the datasets by imputation, yielding a common set of 8671 markers (mostly driven by the most densely genotyped panel LG/JxSM/J F34).

Overall, we observed moderate to little evidence of shared QTLs, even though the panels share a common SM/J parental line (mean Pearson’s correlation = 0.05). In principle, QTL sharing may be due to the same haplotype combinations segregating in separate panels. Alternatively, this could be due to multiple mutations affecting the same genes. Regardless, we found little evidence here to suggest this being a major driver behind the QTL signals. This is in contrast to the comparison between the two LG/JxSM/J F2 panels, whose LOD contour track with each other (Fig. 5, Pearson’s correlation: 0.59, c.f. F2 panels involving MYK: −0.18, −0.14 and −0.08; NZB: −0.08; 0.03 and 0.07). In fact, the MYK x SM/J panel is the only one consistently showing a negative correlation with the other panels. This is underscored by the 6 genome-wide significant *bw16w* QTLs, where only *bw16wq1* seem to be shared broadly with the other F2 panels. More broadly below the significant thresholds, there was a slight overlap for a suggestive QTL on Chr4 with the LG/J panels, but this would have been speculative at best without further relying on the overlap in their corresponding F3 and F34 panels. In a full, combined meta-analysis, we observed a strong QTL on Chr14, but in an intermediate, non-overlapping position between *bw16wq14* and the telomeric QTL from the Palmer LG/JxSM/J F2 panel (Fig. 5).

**Fig. 5.**
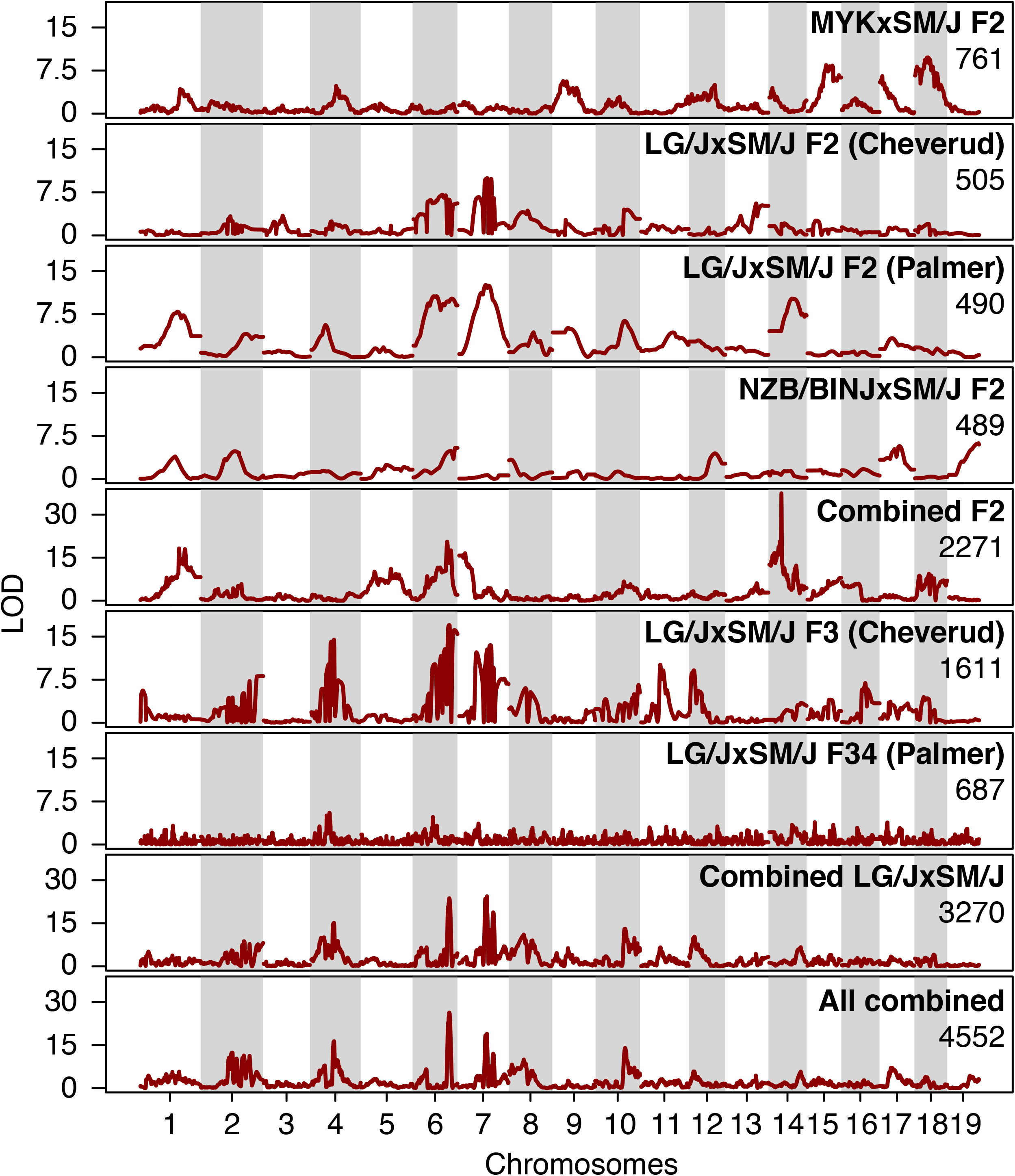
Largely distinct genetic basis of body weight control across a set of crosses sharing an SM/J parent. A meta-analysis was conducted by combining six mapping panels involving the SM/J strain, giving a total sample size of 4552 individuals. The extent of statistical support is shown for each panel, as well as groups of panels together, taking their origins into account. Among the panels, only the MYK x SM/J cross from this study involved wild mice. At the broader QTL level, we did not observe strong overlaps between the Faroese panel and the other panels. Due to the larger size of the various LG/J x SM/J panels, much of the shared signals appear to correspond to panels involving LG/J.

Given the lack of clear overlap across the panels, it was perhaps notable that mapping using all pooled samples seem to show some surprising interactions between the panels. Whereas the QTLs on Chr1 and Chr6 seem to benefit from pooling, the suggestive overlapping QTL on Chr4 now appear to have little to no support. Taken at face value, it would support the interpretation that MYK and LG/J carry distinct alleles on Chr4, despite the appearance of overlapping QTL. On Chr14, there is now a striking QTL that appear to overlap a peak from the Palmer F2 panel, but the signal from other individual F2 panel would appear to be unremarkable. Part of the challenge in interpreting the combined mapping would stem from the mixed coding of all non-SM/J alleles together, which would cause poor fits of the QTL model when in fact multiple allelic effects exist.

Outside of the F2 panels, our results follow broad established patterns: increased genetic resolution is associated with advanced intercrossing like F3 and F34 here, with the latter providing high resolution but relatively decreased mapping power. To be clear, the apparent mapping power in F2 panels is in itself also an artefact due to strong linkage, distorted in both directions from synergistic or opposing effects between linked loci. Combined mapping here shows a clear genetic signal, however, since most of the panels are made up of LG/J x SM/J progenies, the LOD profile from the combined mapping appear to correspond largely to that genetic background (Fig. 5, bottom panels).

Overall, our results show that there may be limited sharing in the genetic basis of body weight changes in the laboratory and in the wild. Whether using a formal QTL analysis like above, or by comparing genomic intervals with the parallel selected regions found across body weight selected laboratory lines (Chan *et al*., 2012), we did not detect a correlation. Additionally, comparison with the previous example of island gigantism on Gough Island also showed almost complete non-overlap in 16 total QTLs, even though the two studies together detected QTLs on 15 out of 20 chromosomes (with the lone exception of the QTLs on Chr9 *bw8wq9, bw14wq9* and *bw16wq9*) (Gray *et al*., 2015). In that sense, the Faroese mouse seem to have largely followed its own unique genetic trajectory, and the laboratory and natural examples of outwardly similar changes represent largely distinct sampling of available genetic variation.

## Discussion

The house mouse on the Faroe Islands is a remarkable example of rapid evolution among wild mice. Since their scientific discovery and description more than a century ago, the Faroese mouse has attracted considerable interest in their large size and their unique phenotypes (Eagle Clarke, 1904; Degerbøl, 1942; Berry *et al*., 1978). Berry and colleagues have first noted a great heterogeneity between the islands in both morphological features and allozyme diversity, and hypothesized that colonization bottlenecks as well as selection combined to cause rapid evolution within a span of only two to three hundred years (Berry *et al*., 1978). Our own genomic survey and population simulation has discovered a much older colonization followed by hybridization between the initial *M. m. domesticus* colonizers and *M. m. musculus* immigrants in the past 300 – 700 years, consistent with the influx of *M. m. musculus* mice following the switch from Viking to Danish control in the Faroe Islands in 1380 (Jones *et al*. in prep.). In this other study, we concluded that the bulk of introgressed *musculus* genome did not behave different from a neutral process, this hybridization event was likely to have contributed to the unique characteristics of the Faroese mice. In the current study, we now try to complete the picture by connecting the unique phenotypes of this population to specific genomic regions.

Here we conducted a genetic mapping experiment between a large island mouse from the Faroe Islands and a small laboratory strain. This mapping panel allowed us to recover a large number of QTLs corresponding to different types of traits, including body weight, length, organ weight and growth hormone levels. The use of a common laboratory small strain makes it possible to directly compare against additional laboratory panels sharing the same strain, and we found that there is only minimal evidence of QTL sharing between these different examples of body weight changes. Likewise, a broader comparison showed that much of the genetic architecture of body weight changes are distinct, either in comparison with laboratory mice or with Gough Island mice. We interpret this to support a highly polygenic genetic architecture for body weight and length in the mouse species complex, as well as the strong sampling effects from inbreeding or colonization.

Among our QTLs, we found a stronger trend towards finding QTLs for later growth. In fact, for our single trait QTL analysis, the one trait with no detected QTL was body weight at 4 weeks, presumably due to a limited sample size (199 measurements, cf. an average of 527 measurements in the rest). This is in contrast to the Gough Island study, where the authors found greater relevance in earlier growth phases (Gray *et al*., 2015). In fact, by partitioning our data according to cross direction, we have found support for a MYK parent-of-origin effect contributing to greater overall growth by week 16, despite starting with slightly lower weaning weights. This genetic signal, however, must be balanced against other possible parent-of-original or maternal effects, which we would not be able to disentangle from our cross design (see Hager *et al*., 2008). Besides other major factors such as having different parental lines and methodologies, the difference in QTL effects on different growth phases may also explain the apparent non-overlap in major QTLs.

Alternatively, the *musculus* hybridization event may have contributed to a unique genetic make-up not observed elsewhere. There is now increasing evidence from many natural systems that such events, while rare, can help create the conditions for novel phenotypes and adaptations and in extreme cases, speciation (Song *et al*., 2011; Sankararaman *et al*., 2014; Huerta-Sánchez *et al*., 2014; Nolte and Tautz, 2010; Mallet, 2007; Heliconius Genome Consortium, 2012). One tentative data-point in support of this would be a somewhat larger effect size at the body weight QTLs than those reported in the Gough Island or even in the LG/J x SM/J crosses (Gray *et al*., 2015; Cheverud, 1996; Norgard *et al*., 2008) (but see QTLs from divergent selection lines, e.g., BEH x BEL) (Brockmann *et al*., 2004). This is despite the larger difference in the body weight between the Gough Island mice and WSB compared to MYK–SM/J here (and accordingly, the MYK allele increases body weight only in six out of ten QTLs). Additionally, we were able to explain a substantial proportion of variation in tail length, a trait known to differ between *musculus* and *domesticus* mice. That said, we emphasize here that the genetic resolution is far below what would be needed to make a formal comparison against introgressed *musculus* loci, and we were unable to detect increased overlap or association with parallel selected loci reported from artificially selected lines (Chan *et al*., 2012). Settling this question would require extensive further genetic work beyond F2 generations, similar to the advanced intercross populations in LG/J x SM/J (Norgard *et al*., 2008; Parker *et al*., 2011).

Unlike our previous study comparing signatures of selection at two specific parallel selected loci, here we found no genome-wide correlation or enrichment between the signature of parallel selection and the LOD profile here. There are a number of relevant considerations here, the main one being the coarse resolution coming from QTL studies vs. the far greater precision from haplotype sharing. A second factor is the use of SM/J as a parental strain here, which was not subjected to divergent selection like those artificially selected lines used in the previous study (MacArthur, 1944; Bünger *et al*., 2001).

## Conclusion

We report here our first study on the genetic basis of the remarkable Faroese house mouse, focusing mainly on body size and weight. Besides their large sizes, these mice were also notable for several unusual morphological characters, especially in the skull. This will be the focus of our next study, in which we employ morphometric techniques to capture such variation. Together with our study of the genomic hybridization history in the Faroese mice (Jones *et al*. in prep.), we have made some progress in understanding the variation among the mice in the Faroe Islands. That said, our work focusing on the Mykines population represents only a small part of the overall picture in the Faroe Islands. Further work, such as admixture mapping from more diverse populations like Sandøy may benefit from the higher genetic resolution beyond those achievable in the F2 cross here.

Besides the Faroe mice, we also expect the current study and others following it to help uncover the genetic underpinning of broader morphological variation in island mouse populations. Indeed, Berry and co-workers have highlighted the broad genetic and phenotypic similarity among Northern Atlantic island mouse populations (Berry *et al*., 1978). Rather than being completely unique in their various characteristics, they stated that it would be more appropriate to describe the Faroese mice as a unique *combination* of characters. Given broad interests in islands in evolutionary biology and the findings already uncovered by this and previous studies on other islands (Gray *et al*., 2015; Parmenter *et al*., 2016), we are hopeful that further work in these unusual mice will continue to yield useful results to improve our understanding on the principles governing novel adaptations.

## Acknowledgements

We thank Felicity Jones for input into experimental design, helpful discussion and improving the manuscript. We thank Emilie Hardouin, Eyðfinn Magnussen, Jens-kjeld Jensen and the Johannessens on Mykines for assistance with field collection. We are grateful for the input from Eleanor Jones and Jeremy Searle, which has greatly facilitated planning. We thank the Tautz, Chan and Jones Labs members for support, insightful scientific discussion and improving the manuscript. We thank Christine Pfeifle, Andy Sörensen and other MausTeam members at the MPI for Evolutionary Biology for animal husbandry. We thank Elke Blohm-Sievers and Heike Harre for phenotyping assistance. We thank Christa Lanz for assistance with high-throughput sequencing; Andre Noll for high-performance computing support; and the MPI Tübingen IT team for computational support. We thank Christine Dreyer and Ralf Sommer for reagents for RAD Sequencing. We are grateful to James Cheverud for sharing his QTL data. We thank Peter Carbonetto for helpful input. Y.F.C. is supported by the Max Planck Society. D.T. is a member of the Max Planck Society.

## Material and Methods

### Field Sampling

House mouse (*Mus musculus spp.*) were sampled with live traps (DeuFa, Neuburg, Germany) on Mykines over successive nights in October 2009. In total, 20 mice were caught and introduced into the Mouse House at the Max Planck Institute for Evolutionary Biology in Plön, Germany. Sampling locations and numbers are shown in Table S1. Due to the small size of Mykines, all sampling locations were less than 1 km apart and were treated as a single locality. The mice were then paired, where possible, within sampling sites to establish the MYK strain.

### Animal Care and Use

All experimental procedures described in this study have been approved by the local competent authorities: the Faroese Food and Veterinary Agency and Ministry for Agriculture, Environment and Rural Area, Schleswig-Holstein, Germany (Permit number 97-8/07).

### Genetic cross and phenotyping

Mice from MYK and SM/J strains bred in our facility under common garden conditions were crossed with each other in reciprocal directions to establish 12 F1 families, which in turn generated 841 F2 mice (Table S2). Mice were weaned at 4 weeks of age. Each F2 mouse was housed singly for 16 weeks, and were weighted biweekly for a total of 7 time-points. At weeks 8 and 16, body and tail length measurements were obtained under anesthesia (week 8) or immediately following sacrifice (week 16). In addition, blood plasma was prepared from freshly drawn blood from the heart and the liver was dissected and weighed. Ear clips were taken for the purpose of DNA extraction.

### Growth curves

In addition to bi-weekly weight measurements, we also estimated parameters related to growth in each mouse using the `grofit` package in R (Kahm *et al*., 2010), which estimated for each mouse the growth rate *μ* and asymptote *A* parameters.

### Growth hormone measurements

Plasma level of growth hormone *Insulin-like growth factor 1* (*Igf1)*, *IGF* binding protein 2 and 3 (*Igfbp2* and *Igfbp3*) were determined using enzyme-linked immunosorbent assay (ELISA) kits (ALPCO, Salem, NH, USA), according to manufacturer’s instructions. Colorimetric reactions were quantified by measuring absorbance at 450 nm using a Tecan Infinite M200 PRO (Tecan AG, Schwerte, Germany) microplate reader equipped with `MAGELLAN 7.0` software. Samples were randomized in their positions, and each sample was measured in duplicates. A standard curve was included in every sample plate and was used to estimate concentration. A subset of samples were measured an additional time to determine repeatability. Repeatability for *Igf1*, *Igfbp2* and *Igfbp3* was determined to be 0.95, 0.94 and 0.78 respectively (*Igf1*, n=43; *Igfbp2*, n=44; *Igfbp3*, n=48, repeatability estimated using the `rpt.aov` function in `rptR` (Stoffel *et al*., 2017).

### Reference genome assembly

All co-ordinates in the mouse genome refer to *Mus musculus* reference `mm10`, which is derived from GRCm38.

### Data and code availability

All raw data and code are deposited at the following repository: https://github.com/evolgenomics/FaroeQTL

### Restriction sites-associated DNA (RAD) sequencing

Given the large size of the mouse genome and the coverage required to confidently call genotypes, we chose to use Restriction sites-associated DNA sequencing (RADseq) to concentrate the sequencing around rare restriction cut-sites. RADseq was performed according to Poland *et al*. 2012, and specifically with the same reagents as used in Witte *et al*., 2015, with the following modifications. Instead of *PstI* (a 6-cutter with the recognition motif C, TGCA’G), *SbfI-*HF (New England Biolabs GmbH), which recognizes the 8-nt motif CC, TGCA’GG but shares the same TGCA 3’ overhang was used together with *MseI*, in order to further enrich the sequencing library for a smaller subset of sites. DNA from the F2 panel of 841 mice were extracted from ear clips. The DNA from each mouse was double digested with *SbfI*-HF and *MseI* and ligated to the adapters. The resulting library was subjected to size selection (400 – 600 bp) using gel electrophoresis. The library was normalized to the same DNA concentration. Sets of 94 libraries were pooled together, amplified by thermocycling using universal primers and sequenced by a HiSeq 2000 (Illumina Inc., San Diego, CA, USA) at the Genome Core Facility at the MPI Tübingen Campus. The overall sequencing output was inspected and about 10% of the samples re-run to ensure sufficient sequence coverage for genotype calling.

### Sequence demultiplexing and genotyping pipeline

The F2 panel was sequenced across a total of 11 HiSeq2000 lanes, including re-runs. In each lane, the sequencing data was pre-processed and demultiplexed using the package `Short Read` (`SHORE`). Briefly, `fastq` files from each lane were demultiplexed into each well via a set of in-line barcodes (5 – 10 nt) in Read1 with the parameter `--barcode-mismatches=1`. In addition to the F2 samples, the two grandparental samples were separately whole-genome shotgun sequenced to approximately 15x coverage by a HiSeq2000 (Illumina) at the Cologne Center for Genomics. Sequenced data were pre-processed using a pipeline consisting of data clean-up, mapping, base-calling and analysis based upon `fastQC v0.10.1` (www.bioinformatics.babraham.ac.uk); `trimmomatic v0.33` (Bolger *et al*., 2014); `bwa v0.7.10-r789` (Li and Durbin, 2010); `GATK v3.4-0-gf196186` modules `BQSR`, `MarkDuplicates`, `IndelRealignment` (McKenna *et al*., 2010; DePristo *et al*., 2011). Genotype calls were made using a pipeline consisting of `samtools mpileup` and `bcftools call` module under the multiallelic mode. The raw genotype calls were filtered using the parameters `TYPE=“snp” && N_SAMPLES > 100 && MAF > 0.25` and only informative positions from the two parents were retained. Using custom `Perl` scripts, we used the two parental lines to polarize the genotypes and averaged the frequency calls over sliding windows of 250 kbp by 50 kbp steps. These form the genotype datasets we used for the subsequent linkage mapping step.

### Linkage mapping

Linkage mapping was performed in R using the packages R/qtl (Broman *et al*., 2003) and R/qtlRel (Cheng *et al*., 2010). Gentoype data was coded as “M” for Mykines, “H” for heterozygous and “S” for SM/J. Due to the use of the Faroe mouse parental line, we rebuild a genetic map from this dataset using the Kosambi map function. The resulting map has a total length of 2379 cM (65 – 200 cM). In total, 22 phenotypes were retained for this analysis, consisting of weight, length and plasma protein measurements, as well as a set of covariates such as sex, family history, age and cross directions. We applied Box–Cox transformation to numeric datasets, resulting in z-standardized, mean-centered phenotypes with improved normality. Genetic mapping was performed on the transformed dataset. We also obtained major axes of variation through principal component analysis. For each trait, we applied corrections for family, sex and cross directions following a backwards model selection procedure in which we simplify from a full additive and interactive set of covariates to the minimal set based on the Aikake Information Criterion (AIC). Genome-wide significance thresholds were determined from 1000 permutations. For QTL mapping with relatedness correction under `QTLrel`, relatedness was estimated from marker genotypes at non-focal chromosomes and was fitted as a random effect. Following QTL detection, the effect sizes were estimated by fitting a QTL model against the original, untransformed phenotypes.

### Meta-analysis

The analysis across different mapping panel was performed used QTLrel (Cheng *et al*., 2010), following closely the procedure described in Parker et al., 2014 (Parker *et al*., 2014). Briefly, phenotype and genotype data from the Cheverud and Palmer labs, as well as from Stylianou et al., 2006 were obtained (Cheverud, 1996; Stylianou *et al*., 2006; Norgard *et al*., 2008; Parker *et al*., 2014). The genotype data was coded as the number of SM/J allele in each panel. Then QTL mapping with relatedness correction was performed, using marker-caluclated measures of relatedness. Since our main focus here is the broader comparison of mapping results, we did not attempt extensive bootstrapping analyses to determine the significant threshold in the combined analysis.

## Supplemental Materials

### Supplemental Figures

**Figure S1.**
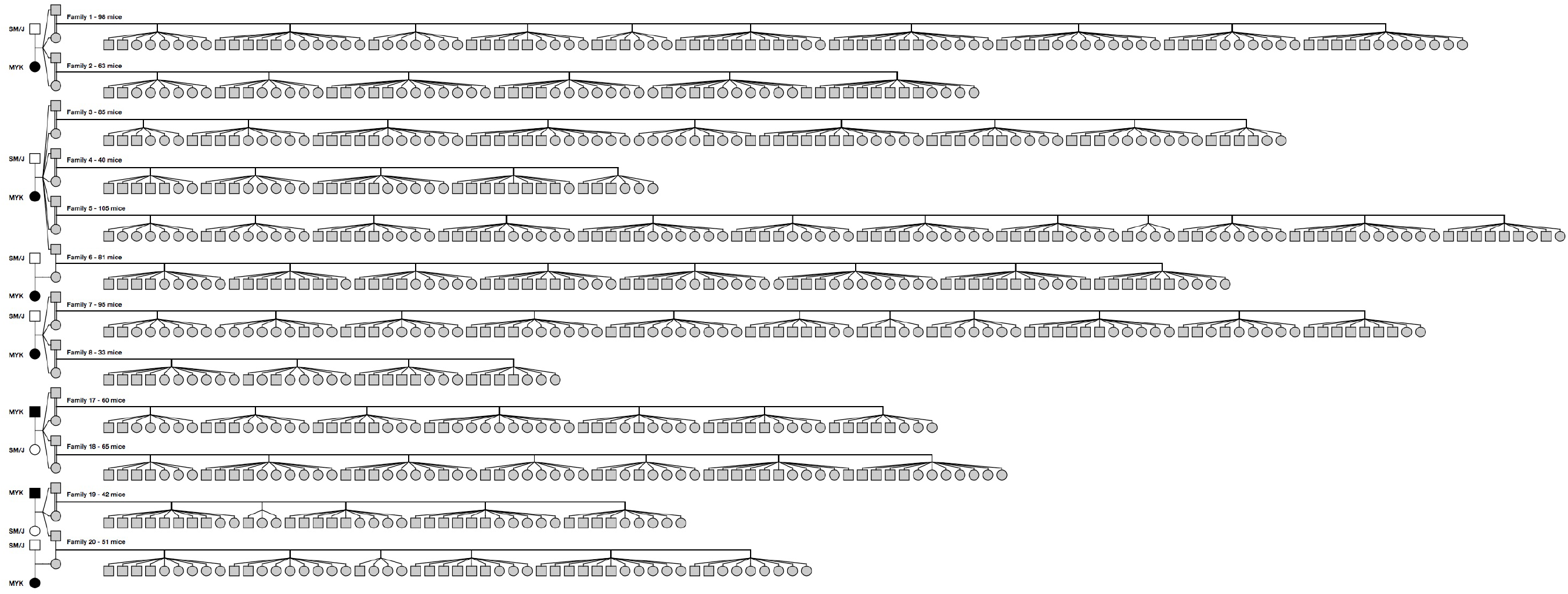
Pedigree of the MYK x SM/J cross. Shown are the pedigree relationship between the animals in the current study (males: squares; females: circles; MYK: black; SM/J: white), with the F2 families rotated to fit onto the page. Each horizontal row represent the offspring from an F2 family, with litters indicated by groups.

**Figure S2.**
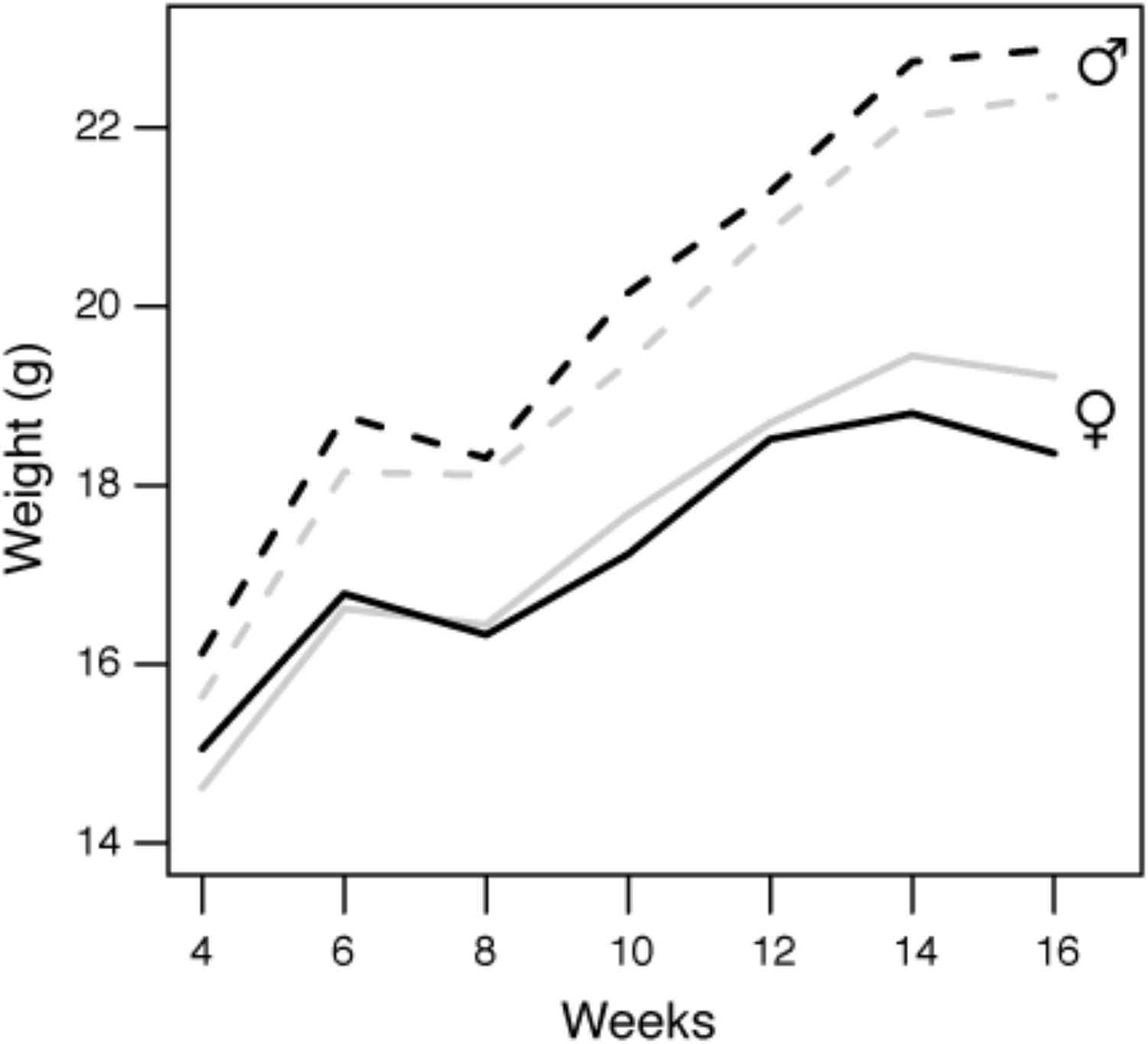
Parent-of-origin effect on weight gain. Shown here are the mean growth curve of the F2 mice, split by cross direction. The main difference in cross direction can be best shown by tracking the X chromosome: F2 females all carry an X chromosome from the paternal grandmother due to the pedigree structure. Here, we observed a significant interaction with cross direction in females (solid lines; repeated measures ANOVA, *P <* 0.0003; but *P* < 0.89 in males, dashed lines). F2 females carrying an MYK X chromosome were born with lower weaning weight at 4 weeks of age, but gained greater weight by week 16 There was no discernible effect among males.

### Supplemental Tables

**Table. S1.**
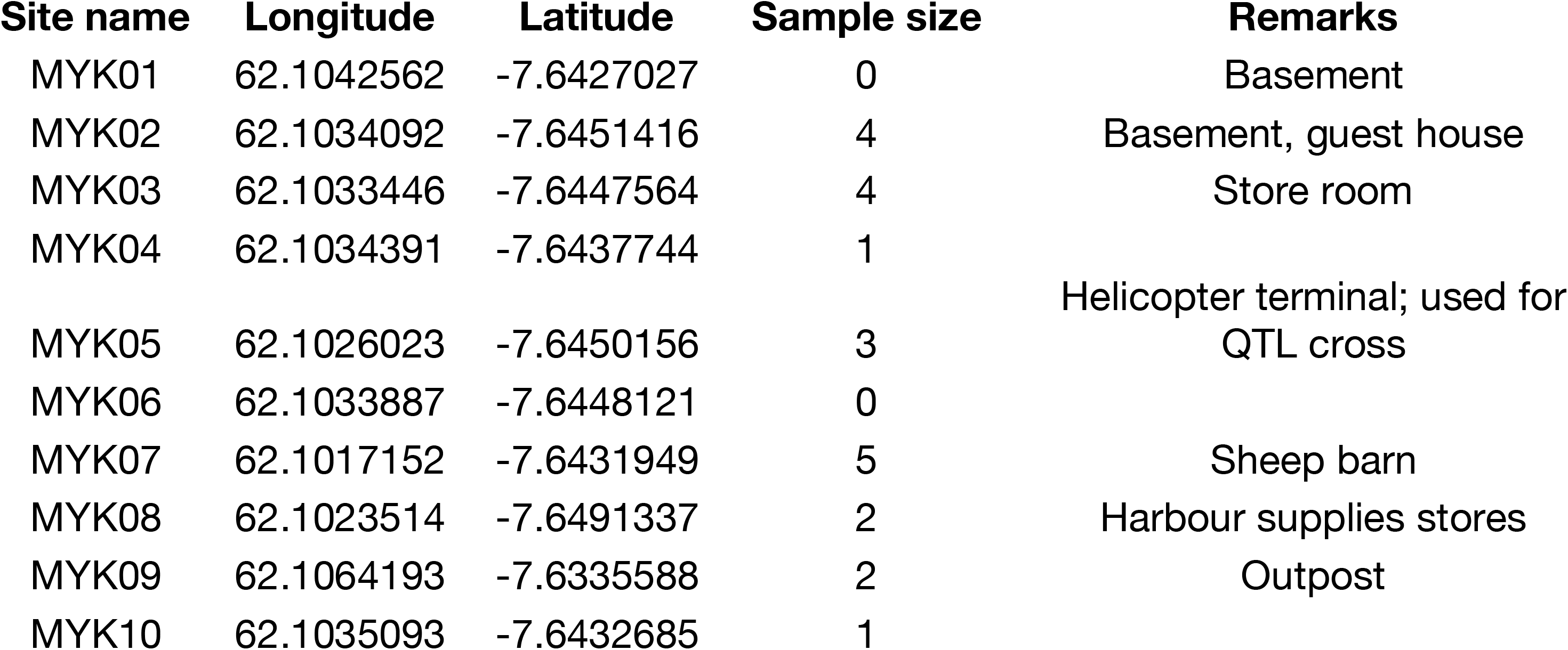
Sampling locations.

**Table. S2.**
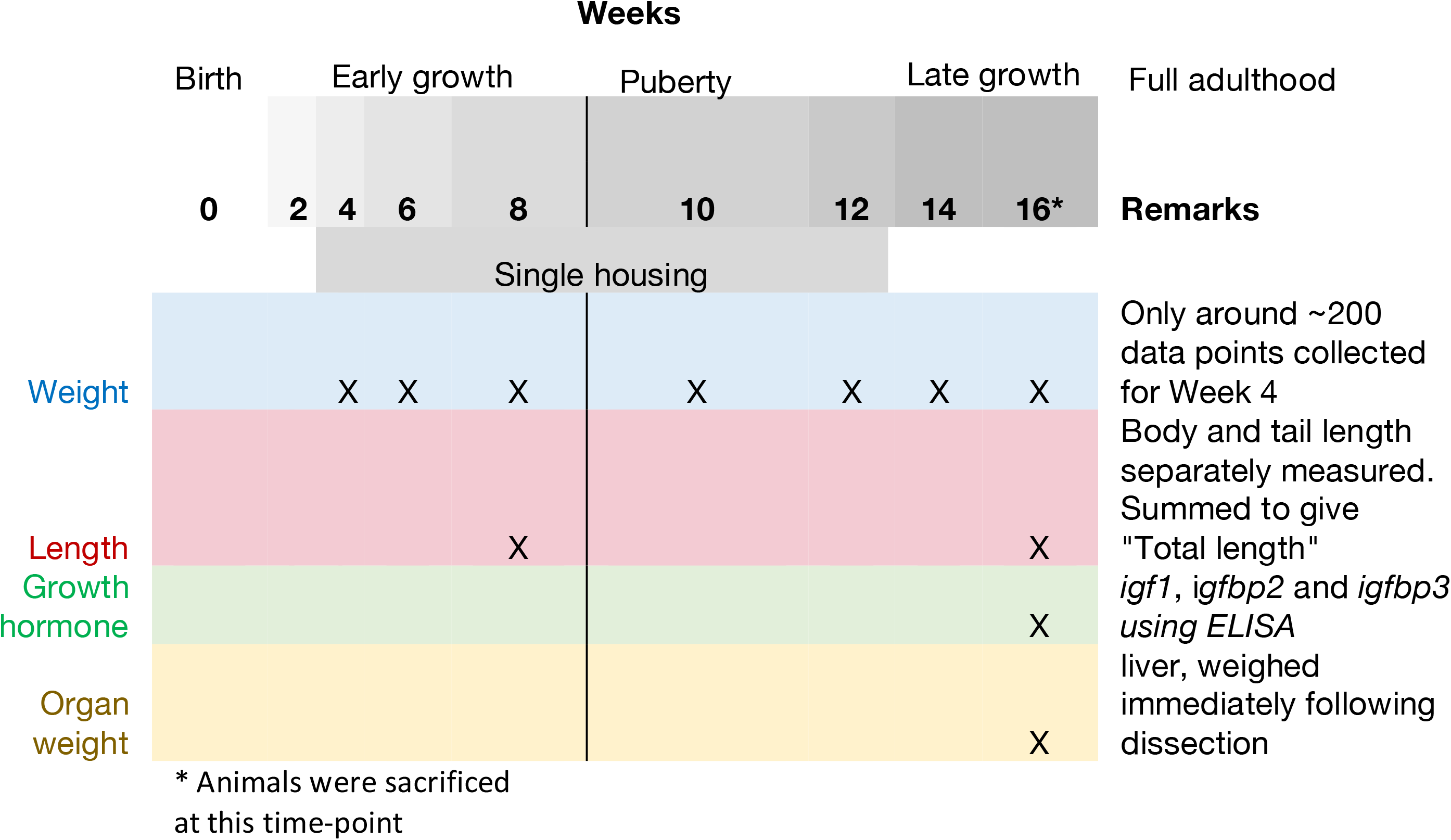
Phenotyping scheme. This table schematically represents the phenotyping time-course of each F2 mouse in the mapping experiment.

**Table S3.** Correlation matrix between measured traits.

|  | IGF1_ELISA | IGFBP2_ELISA | IGFBP3_ELISA | GrowthModel_A | GrowthModel_μ | BodyWeight.4Wk | BodyWeight.6Wk | BodyWeight.8Wk | BodyWeight.10Wk | BodyWeight.12Wk | BodyWeight.14Wk | BodyWeight.16Wk | LiverWeight.16Wk | TotalLength.8Wk | TotalLength.16Wk | TailLength.8Wk | TailLength.16Wk | BodyLength.8Wk | BodyLength.16Wk |
| --- | --- | --- | --- | --- | --- | --- | --- | --- | --- | --- | --- | --- | --- | --- | --- | --- | --- | --- | --- |
| IGF1_ELISA | 1.00 | 0.39 | 0.71 | 0.52 | 0.09 | 0.13 | 0.31 | 0.43 | 0.41 | 0.47 | 0.55 | 0.61 | 0.58 | 0.35 | 0.51 | 0.25 | 0.34 | 0.34 | 0.55 |
| IGFBP2_ELISA | 0.39 | 1.00 | 0.38 | 0.31 | 0.12 | -0.04 | 0.14 | 0.25 | 0.23 | 0.28 | 0.35 | 0.38 | 0.36 | 0.19 | 0.30 | 0.06 | 0.21 | 0.25 | 0.32 |
| IGFBP3_ELISA | 0.71 | 0.38 | 1.00 | 0.36 | 0.11 | 0.05 | 0.23 | 0.32 | 0.35 | 0.39 | 0.46 | 0.51 | 0.57 | 0.26 | 0.39 | 0.18 | 0.23 | 0.25 | 0.46 |
| GrowthModel_A | 0.52 | 0.31 | 0.36 | 1.00 | -0.04 | 0.60 | 0.56 | 0.71 | 0.70 | 0.84 | 0.89 | 0.92 | 0.70 | 0.58 | 0.69 | 0.45 | 0.56 | 0.53 | 0.67 |
| GrowthModel_μ | 0.09 | 0.12 | 0.11 | -0.04 | 1.00 | -0.26 | 0.03 | 0.37 | 0.37 | 0.24 | 0.22 | 0.20 | 0.21 | 0.24 | 0.24 | 0.19 | 0.21 | 0.21 | 0.21 |
| BodyWeight.4Wk | 0.13 | -0.04 | 0.05 | 0.60 | -0.26 | 1.00 | 0.52 | 0.49 | 0.45 | 0.49 | 0.42 | 0.40 | 0.22 | 0.47 | 0.38 | 0.50 | 0.36 | 0.31 | 0.34 |
| BodyWeight.6Wk | 0.31 | 0.14 | 0.23 | 0.56 | 0.03 | 0.52 | 1.00 | 0.70 | 0.64 | 0.60 | 0.60 | 0.58 | 0.47 | 0.55 | 0.52 | 0.44 | 0.42 | 0.49 | 0.50 |
| BodyWeight.8Wk | 0.43 | 0.25 | 0.32 | 0.71 | 0.37 | 0.49 | 0.70 | 1.00 | 0.85 | 0.82 | 0.82 | 0.74 | 0.60 | 0.72 | 0.65 | 0.54 | 0.52 | 0.68 | 0.63 |
| BodyWeight.10Wk | 0.41 | 0.23 | 0.35 | 0.70 | 0.37 | 0.45 | 0.64 | 0.85 | 1.00 | 0.81 | 0.80 | 0.72 | 0.60 | 0.63 | 0.63 | 0.48 | 0.49 | 0.57 | 0.61 |
| BodyWeight.12Wk | 0.47 | 0.28 | 0.39 | 0.84 | 0.24 | 0.49 | 0.60 | 0.82 | 0.81 | 1.00 | 0.87 | 0.78 | 0.65 | 0.61 | 0.65 | 0.45 | 0.49 | 0.58 | 0.66 |
| BodyWeight.14Wk | 0.55 | 0.35 | 0.46 | 0.89 | 0.22 | 0.42 | 0.60 | 0.82 | 0.80 | 0.87 | 1.00 | 0.91 | 0.73 | 0.64 | 0.73 | 0.45 | 0.55 | 0.62 | 0.74 |
| BodyWeight.16Wk | 0.61 | 0.38 | 0.51 | 0.92 | 0.20 | 0.40 | 0.58 | 0.74 | 0.72 | 0.78 | 0.91 | 1.00 | 0.80 | 0.60 | 0.73 | 0.42 | 0.55 | 0.58 | 0.74 |
| LiverWeight.16Wk | 0.58 | 0.36 | 0.57 | 0.70 | 0.21 | 0.22 | 0.47 | 0.60 | 0.60 | 0.65 | 0.73 | 0.80 | 1.00 | 0.44 | 0.61 | 0.32 | 0.42 | 0.41 | 0.64 |
| TotalLength.8Wk | 0.35 | 0.19 | 0.26 | 0.58 | 0.24 | 0.47 | 0.55 | 0.72 | 0.63 | 0.61 | 0.64 | 0.60 | 0.44 | 1.00 | 0.79 | 0.82 | 0.77 | 0.85 | 0.60 |
| TotalLength.16Wk | 0.51 | 0.30 | 0.39 | 0.69 | 0.24 | 0.38 | 0.52 | 0.65 | 0.63 | 0.65 | 0.73 | 0.73 | 0.61 | 0.79 | 1.00 | 0.73 | 0.89 | 0.60 | 0.86 |
| TailLength.8Wk | 0.25 | 0.06 | 0.18 | 0.45 | 0.19 | 0.50 | 0.44 | 0.54 | 0.48 | 0.45 | 0.45 | 0.42 | 0.32 | 0.82 | 0.73 | 1.00 | 0.84 | 0.41 | 0.43 |
| TailLength.16Wk | 0.34 | 0.21 | 0.23 | 0.56 | 0.21 | 0.36 | 0.42 | 0.52 | 0.49 | 0.49 | 0.55 | 0.55 | 0.42 | 0.77 | 0.89 | 0.84 | 1.00 | 0.48 | 0.53 |
| BodyLength.8Wk | 0.34 | 0.25 | 0.25 | 0.53 | 0.21 | 0.31 | 0.49 | 0.68 | 0.57 | 0.58 | 0.62 | 0.58 | 0.41 | 0.85 | 0.60 | 0.41 | 0.48 | 1.00 | 0.57 |
| BodyLength.16Wk | 0.55 | 0.32 | 0.46 | 0.67 | 0.21 | 0.34 | 0.50 | 0.63 | 0.61 | 0.66 | 0.74 | 0.74 | 0.64 | 0.60 | 0.86 | 0.43 | 0.53 | 0.57 | 1.00 |

**Table S4.** Datasets used in the SM/J meta-analysis.

| **Panel** | **Parent 1** | **Parent 2** | **Generation** | **N\*** | **Reference** |
| --- | --- | --- | --- | --- | --- |
| Chan F2 | MYK | SM/J | F2 | 761 | This study |
| Cheverud F2 | LG/J | SM/J | F2 | 505 | Cheverud *et al.*, 1996 |
| Cheverud F3 | LG/J | SM/J | F3 | 1611 | Norgard *et al.*, 2008 |
| Palmer F2 | LG/J | SM/J | F2 | 490 | Parker *et al.*, 2011 |
| Palmer F34 | LG/J | SM/J | F34 | 687 | Parker *et al.*, 2011 |
| Stylianou F2 | NSB/BINJ | SM/J | F2 | 489 | Stylianou *et al.*, 2006 |

\* retained samples with comparable phenotypes

